# Time-course of antipredator behavioral changes induced by the helminth *Pomphorhynchus laevis* in its intermediate host *Gammarus pulex*: the switch in manipulation according to parasite developmental stage differs between behaviors

**DOI:** 10.1101/2023.04.25.538244

**Authors:** Thierry Rigaud, Aude Balourdet, Alexandre Bauer

## Abstract

Many trophically transmitted parasites with complex life cycles manipulate their intermediate host’s antipredatory defenses in ways facilitating their transmission to final host by predation. Some parasites also protect the intermediate host from predation when noninfective during its ontogeny. The acanthocephalan *Pomphorynchus laevis*, a fish intestinal helminth, infecting freshwater gammarid amphipods as intermediate hosts, is using such a strategy of protection-then-exposure to predation by the definitive host. However, the whole time-course of this sequence of behavioral switch is not yet known, and only one antipredator behavior has been studied to date. Here we show that the protective part of this manipulation begins quite late during the parasite ontogeny, suggesting that the advantages overpass the costs induced by this protective manipulation only at this stage. We confirmed that the refuge use behavior is showing a switch in the few days following the stage mature for the definitive host (switching from overuse to underuse). However, such a switch was not observed for the gammarids activity rate, a behavior also known to make the host less conspicuous to predators when weak. While we predicted a low activity during early development stages, then a switch to high activity, we observed general decrease in activity during the parasite ontogeny. The possible causes for this discrepancy between behaviors are discussed. All these behavioral changes were observed mostly when animals were tested in water scented by potential predators (here brown trout), suggesting condition-dependent manipulation.

## Introduction

Heteroxenous parasites require successive hosts to complete their life cycle (Poulin, 2007). A successful transfer from one host to another is critical to the fitness of such parasites, and numerous parasite species rely on trophical transmission to achieve this goal. In this process, downstream infected host(s) – the intermediate host(s) – must be consumed by the upstream host – the definitive host – where the parasite reproduce. Parasite-induced phenotypic alterations (PIPA) may facilitate the completion of the parasite’s life cycle (Moore, 2002; Poulin, 2010). Alterations of host behavior are particularly frequent in trophically-transmitted parasites, with infected hosts showing behaviors increasing their chances to be predated by their parasite’s final host e.g. (Bethel & Holmes, 1977; Moore, 1983; Lagrue et al., 2007), reviewed, for example, in Poulin (2010). These parasites generally alter the antipredator behaviors of their intermediate hosts (Svensson et al., 2022), and some parasites are even able to reverse their intermediate host’s fear of predator cues (Berdoy et al., 2000; Perrot-Minnot et al., 2007). These host behavioral changes favoring parasite transmission are adaptive for parasites but detrimental to the host. They are therefore seen as an extended phenotype of parasites and are referred to as parasitic manipulation (Hughes et al., 2012).

The understanding of the adaptive significance of PIPA was reinforced by studies highlighting variations in manipulations depending on the ontogeny of the parasite. Parker et al. (2009) predicted that the trophically-transmitted parasites showing the strongest selective advantage would be those able to protect from predation when not infective to the definitive host and then switch to an increase of exposure to predation when infectious. This phenomenon might have been selected because trophic transmission occurring before the parasite has reached the infective stage would result in zero fitness (Parker et al., 2009). Ideally, then, an optimal temporal sequence of manipulation could consist of a ‘protect-then-expose’ strategy against predation. Accordingly, reinforced antipredatory behaviors were observed during the early stages of the ontogeny in a number of parasite species (e.g. Hammerschmidt et al., 2009; Médoc & Beisel, 2011; Bailly et al., 2018) and direct testing for predation rate of uninfected versus infected hosts confirmed the existence of this switch in predation rates (Dianne et al., 2011; Weinreich et al., 2013). This phenomenon has been studied in detail in a few model organisms only. An iconic model is the tapeworm *Schistocephalus solidus*, where infection in its copepod first intermediate host firstly reduces the host activity and recovery time after a disturbance mimicking a predator attack (Hammerschmidt et al., 2009), resulting in a decrease in predation (Weinreich et al., 2013). This manipulation occurs rapidly after the establishment of the infection within the copepod body (Benesh, 2019). After infectivity for the next host is reached, the behavioral pattern reverses, infected copepods becoming more active and conspicuous to predators (Hammerschmidt et al., 2009). In this system, the intensity of host manipulation varied mainly according to parasite populations, and responded to selection on parasites, confirming that this trait can be a parasite extended phenotype (Hafer, 2018; Hafer-Hahmann, 2019), rather than a trait of the host also favoring the parasite fitness (Jensen et al. 2023).

Another well-studied parasite-host system for switching in behavioral parasitic manipulation is the acanthocephalan *Pomphorhynchus laevis* infecting the amphipod *Gammarus pulex*. Here, the infection by the non-infective stage (the acanthella) protects the host from predation by increasing the rate of refuge use in the amphipod (Dianne et al., 2011), a trait protecting gammarids from predation by fish (Abjornsson et al., 2004). A switch then occurs few days after the parasite reached the cystacanth stage (infective for the next host), exposing the infected host to fish predation (Dianne et al., 2011). In this system, no selection was made on parasites to confirm this extended phenotype, but the intensity of manipulation depends on the parasite strains (Franceschi, Bollache, et al., 2010) or populations (Franceschi, Cornet, et al., 2010; Labaude et al., 2015), suggesting that manipulation is a parasite trait. However, here, a number of data are still lacking to further understand the ‘protect-then-expose’ strategy. For example, it is not known if *P. laevis* invests in host protection early in its ontogeny, knowing that the development of the acanthella stage necessitates quite a long time, i.e. between 2 and 3 months depending on temperature (Labaude et al., 2020). No data are available on the course of behavioral changes during the whole duration of the acanthellae stage. Indeed, measures of refuge use or predation rates were made at the end of this acanthella stage, few days before the parasite reached the infective stage (Dianne et al., 2011, 2014). However, a long time spent in a host would necessitate a long investment in protection. Dianne et al. (2014) identified a cost of the protective manipulation in terms of host food intake. Such a cost, if imposed on a long time, could impact the survival of the host, therefore limiting a long protective manipulation during parasite ontogeny. The life-cycle of *Schistocephalus solidus*, much shorter (around two weeks before being infective in the copepod host), probably allows early investment in the protective manipulation (Hammerschmidt et al., 2009). In addition, it is known that manipulation of gammarids by *P. laevis* is multidimentional, i.e. that multiple traits are changed in the hosts by the parasite infection (Perrot-Minnot et al., 2014; Fayard et al., 2020). However, we still do not know if multiple antipredatory traits are changed during protective manipulation. The activity rate, for example, known to be reduced as an antipredator defense in gammarids (Williams & Moore, 1985; Bollache et al., 2006) should be reduced during the acanthella stage, and then switch to an increase at the cystacanth stage, but this has never been tested. Finally, while Dianne et al. (2011) found that acanthellae induced an increased use of refuges only in the presence of predator cues (fish scent), Dianne et al. (2014) found contradictorily that this protective manipulation in *P. laevis* was observed both in presence or absence of predatory cues, suggesting that the protective manipulation is not condition-dependent. However, again, no data is available on the entire ontogeny of the acanthella.

In this study, we analyzed the behavioral changes induced by *P. laevis* in *G. pulex*, for the first time during the whole ontogeny of parasites. Two anti-predatory behaviors, namely the intensity of host refuge use and the host activity, were measured after experimental infections (as recommended by Poulin & Maure, 2015) and under two conditions: with or without predation cues. Three main questions were addressed: (i) is the protective manipulation present during early acanthella stages of parasite’s larval development? (ii) is this manipulation condition(predator)-dependent during the whole larval ontogeny? (iii) are these two different anti-predatory behaviors affected in the same way by *P. laevis* during its ontogeny?

## Material and Methods

### Animals and experimental infections

Collection, maintenance and infection procedure were as described in Franceschi et al. (2008). In brief, uninfected *G. pulex* amphipods were collected in February 2012 from a small tributary of the Suzon River, Burgundy, eastern France (47°23’56.19’’N, 4°50’31.13’’E). Only male gammarids were kept for the experiment to avoid any interference of the female reproductive cycle on the outcome of parasite infection (Franceschi *et al*. 2008). They were acclimated in the laboratory for 3 weeks before experimental infections under a 12:12 light:dark cycle and fed during the experiment with conditioned elm leaves (*Ulmus laevis*). Adult *P. laevis* parasites were removed from the intestine of naturally infected chub (*Squalius cephalus*) caught in February 2012 in the Vouge River, France (47°09’34.36’’N, 5°09’02.50’’E) and characterized molecularly as described in Franceschi et al. (2008). Parasite eggs were collected from six females of *P. laevis* sampled from four different fish. Male gammarids were allowed to feed for 48 h on a 1 cm^2^ piece of elm leaf, on which about 200 eggs were deposited. A total of 190 *G. pulex* males were exposed to parasite eggs. After exposure, gammarids were maintained in isolation in cups (6 cm diameter) filled with c.a. 50 mL of aerated water. Dead elm leaves were provided as food *ad libitum* and water was changed every week. Twenty control, unexposed, gammarids were handled and maintained under the same conditions as the exposed gammarids.

### Behavioral measurements and survival

Survival was controlled every day post-exposure, and dead gammarids were dissected within the 24 hours post-death in order to control for parasite presence. Due to small size of acanthella, we were unable to detect this presence before 21 days in the 14 *G. pulex* that died (personal observations). Since we were unable to assign hosts to the infected or uninfected categories, the comparisons of refuge use and survival were made after the 21^th^ day post-infection. The experiment was stopped after 83 days post-exposure.

We measured the rate of refuge use every week. Gammarids were introduced individually into a 10.5 × 16 cm rectangular box filled with 330 mL of either control or fish-scented water, with each gammarid kept in the same water type during the whole experiment. Ten of the 20 unexposed control gammarids were tested under fish-scented water, 10 in control water. To produce predator cues, six young brown trouts (*Salmo trutta*), each ∼ 20 cm long, were kept in the laboratory at 15 ± 1 °C under a 12:12 light:dark cycle, in a tank filled with 80 L of tap water previously dechlorinated, oxygenated and UV-treated. Trouts were acclimated for 2 weeks prior to the experiments and fed with a mix of frozen chironomid larvae and alive gammarids in order to strengthen the predation signal (Wudkevich et al., 1997). The water from a similar tank, treated as previously described but without trout scent, was used as a control water for behavioral experiments. Individuals were randomly assigned to these water treatments. An opaque refuge, consisting of half a terracotta saucer (8.5 diameter) in which a 1 cm^2^ opening was made, was placed at one end of each aquarium, covering approximately 18% of the total aquarium area. Refuge use was recorded after an acclimation period of 30 min. The gammarid’s position in the box was then checked every 5 min over a period of 120 min. A score of 1 was given if the gammarid was inside the refuge and a score of 0 if it was found outside. For each gammarid, the total score of refuge use therefore ranged between 0 (gammarid always outside the refuge) and 24 (gammarid always inside the refuge). After behavioral measure, the gammarid returned to its cup, where new aerated water (without fish scent) and food were provided.

The activity of gammarids was recorded the day after the refuge use measurement, using a ViewPoint device and software (©Viewpoint Life Sciences, Inc. – 2010, France). Gammarids were placed individually in Petri dishes (diameter 8.5 cm, height 1 cm) filled with either control water or scented water, this water treatment being the same as for refuge use measurements for each gammarid. After 2 min of acclimation, gammarid activity was recorded with an infrared camera for 5 min (continuous recording), as the proportion of time spent inactive. Inactivity was defined as movements at a speed below 10 mm s^-1^. This speed threshold was determined based on preliminary tests, showing that below 10 mm s^-1^ discrimination is not possible between gammarids moving at very low speed (crawling) and gammarids moving their pleopods for respiration. The 20 control unexposed gammarids were not measured for this trait.

The infection procedure allowed synchronized development of acanthellae. Around 55 days post-exposure, the presence of *P. laevis* acanthellae was unambiguously detectable through the host cuticle (translucent light-orange and shapeless larval stages). Therefore, at that time, all gammarids were assigned to exposed-infected or exposed-uninfected categories (either dead gammarids showed an infection after dissection, or the infection became detectable in living gammarids). It means that behavioral tests were made in blind for the infection status before 55 days post-exposure. The cystacanth stage was reached ten days later (appearing through the cuticle as small bright orange opaque balls).

### Statistical analysis

The survival of gammarids was analyzed using Cox regressions, where the effects of infection status (in a first analysis: unexposed vs exposed-uninfected vs exposed-infected individuals; in a second analysis: uninfected vs infected), water type (scented vs control) and their interaction were analyzed using the ‘survival’ R package (V 3.2). Similar procedures were used in infected individuals only to test the effect of the number of parasites on survival.

Scores for refuge use and activity rate were analyzed as repeated measures using the ‘nparLD’ R software package (V 2.1), with the F2 LD F1 model. This function enables nonparametric analyses of right-censored longitudinal data, allowing the decrease in sample size along time, due to individuals’ death (Noguchi et al., 2012). The effects of water type (scented vs control), infection status (uninfected vs infected) and their interaction were investigated along time, considered here as an ordinal variable. ANOVA-type statistics (AT Statistics), as recommended for small sample sizes (n < 8) were performed (Noguchi et al., 2012), the statistical analysis of time-series with nparLD being based on rank-order of the observed data (see Noguchi et al., 2012 for details). We also report Cliff’s δ with their 95% confidence intervals (Cliff, 1993) as measures of non parametric effect size for the difference in refuge scores or activity rate between infected vs uninfected gammarids at each time step, for each water type. This parameter measures the frequency of values in a given distribution that are larger than the values in another distribution. The values of δ ≥ 0.11 are considered as small, those ≥ 0.28 as medium, and those ≥ 0.43 as large (Kraemer & Kupfer, 2006).

All statistical analyses were performed using R version 4.1.0 software (R Foundation for Statistical Computing, Vienna, Austria).

## Results

We were able to detect unambiguously infections through the cuticle of the hosts at the late acanthella stage around the 55^th^ day post-exposure. Before this stage, parasite infections were only detectable after dissection of dead individuals from day 21. Out of the 176 surviving individuals exposed to parasite eggs, 132 were finally infected (75%). Among them, 60 (45.5%) were infected by only one parasite, 39 (29.5%) were infected by two parasites and 33 (25.0%) were infected by more than two parasites.

The experiment was stopped after 83 days post-infection and all surviving animals were dissected to confirm their infection status and count the parasites. Exposed-uninfected animals did not differ significantly from control unexposed gammarids in either survival or shelter use (supplementary informations). Therefore, in order to increase the power of the behavioral analyses at the end of the experiment, we merged unexposed and exposed-uninfected animals to create the uninfected or not infected category (terminology used thereafter).

The analysis of survival did not fulfill the conditions of proportional hazards throughout the whole period (χ^2^ = 20.9775; 3 d.f.; p = 0.0001). We therefore ran two analyses according to the two main stages of the larval developmental cycle, first during the acanthellae stage, then after the cystacanth stage has been reached. Analyses of the survival during these two periods fulfilled the conditions of proportional hazards (χ^2^ = 2.67542; 3 d.f.; p = 0.44 and χ^2^ = 3.9257; 3 d.f.; p = 0.27, respectively).

Animals survived equally through the acanthella stage regardless of their infection status, but the uninfected individuals survived better than the infected ones after the parasites reached the cystacanth stage (Table 1, Figure 1). The water condition had no significant effect on survival (Table1).

**Table 1:** Effect of water type, infection status by *Pomphorhynchus laevis*, and their interaction on the survival of *Gammarus pulex* individuals, during the acanthella stage development (a) and after the cystacanth stage has been reached (b).

| Source | d.f. | z | p-value |
| --- | --- | --- | --- |
| (a) |  |  |  |
| Water type | 1 | 1.840 | 0.0658 |
| Infection | 1 | 1.039 | 0.2988 |
| Water type*infection | 1 | -1.486 | 0.1374 |
| (b) |  |  |  |
| Water type | 1 | 1.433 | 0.1517 |
| **Infection** | **1** | **-2.562** | **0.0104** |
| Water type*infection | 1 | -1.141 | 0.2540 |

**Figure 1:**
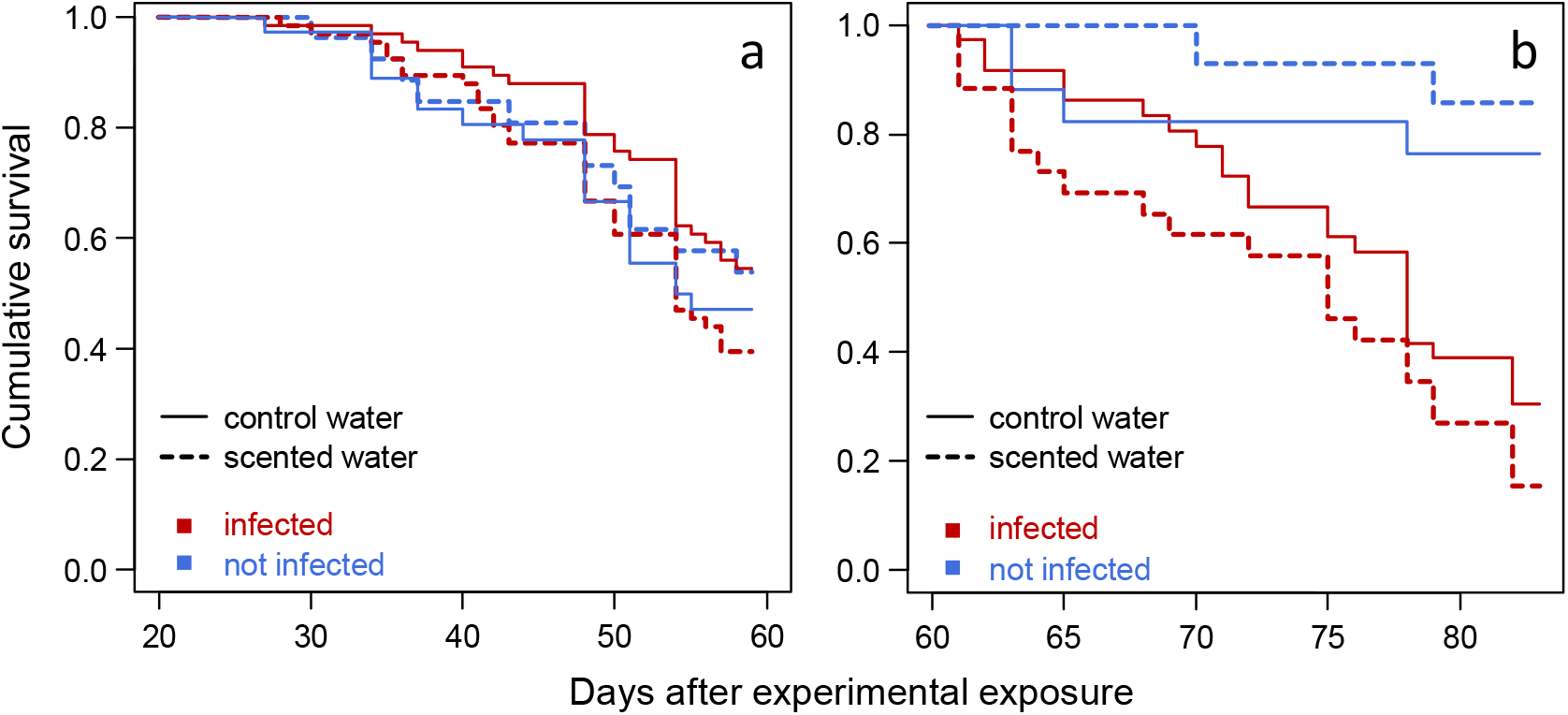
Cumulative survival of *Gammarus pulex* after experimental exposure to *Pomphorhynchus laevis* according to their infection status, during the acanthella stage development (a) and after the cystacanth stage has been reached (b). Animals were maintained in water that was (or not) signed with fish scent. After cystacanth stage was reached, infected *G*.*pulex* suffer higher mortality.

Among the infected individuals, to analyze the effect of the infection intensity on survival, the number of parasites was categorized as “1 parasite” or “more than 1 parasite”. The number of individuals infected by one parasite were 29 in control water and 31 in scented water, while 37 individuals with more than one parasite were found in control water and 35 in scented water. Neither the parasite intensity nor the water type influenced significantly the survival (z = -1.556, p = 0.120 and z = 1.051, p = 0.293, respectively). The interaction was also non significant (z = 1.149, p =0.251).

The refuge use by gammarids varied considerably with time (Figure 2, Table 2), being most of the time moderate at the beginning of our survey, becoming progressively more intensive until reaching high values around 30 to 40 days.

**Figure 2:**
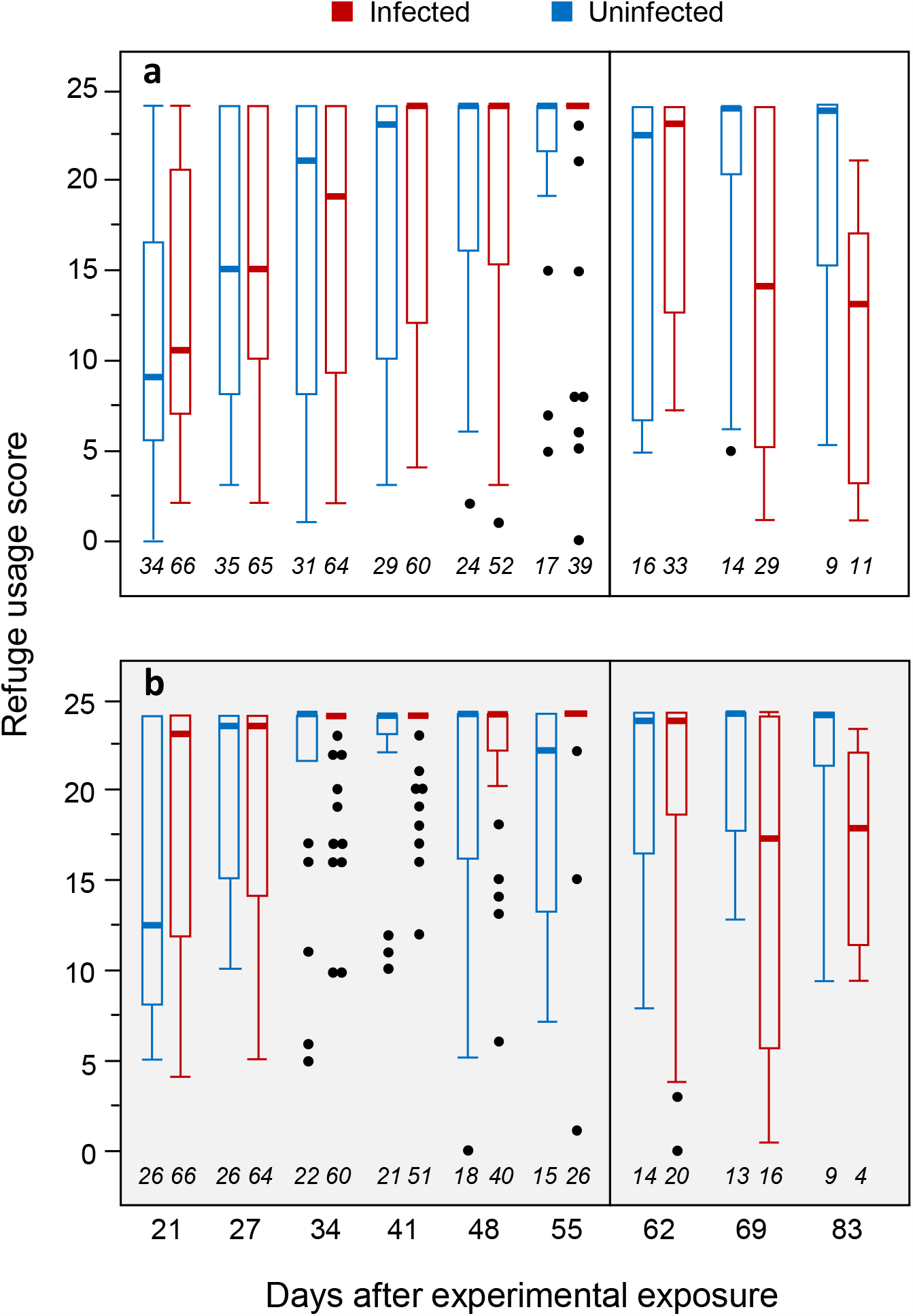
Scores of refuge use by gammarids, according to their infection status by *Pomphorhynchus laevis*, between day 21 and day 83 post-experimental exposure to parasite eggs. Gammarids were tested in control water (a), or water signed with fish scent (b). Scores range from 0 (individuals always outside the refuge) to 24 (individuals always inside the refuge). Thick lines are the medians, boxes are the upper and lower quartiles, whiskers are the upper and lower 1.5*interquartile range, and dots represent outliers. The vertical black line denotes the transition between acanthella and cystacanth stages. Sample sizes are given in italics below each plot. Refuge use increase during acanthealla stages (this phenomenon was stronger under signed water) and then decreased after cyctacanth stage is reached.

**Table 2:** Results of the model from the nparLD analysis, testing for the effects of water type (scented vs. control), infection status (infected with *P. laevis* vs. uninfected) and date of measurement on the scores of refuge use by *G. pulex* individuals.

|  | AT statistics | df | P-value |
| --- | --- | --- | --- |
| **Water type** | **7.8568** | **1** | **0.0051** |
| Infection | 0.9645 | 1 | 0.3261 |
| **date** | **12.9330** | **5.7522** | **< 0.00001** |
| Water type * Infection | 0.4299 | 1 | 0.5120 |
| **Water type \* date** | **2.9745** | **5.7522** | **0.0075** |
| **Infection \* date** | **11.1301** | **5.7522** | **< 0.00001** |
| Water type * Infection * date | 7.5111 | 5.7522 | 0.6030 |

This phenomenon was found regardless of the infection status and the water type, but was found to be more rapid under scented water than control water (Figure 2), the interaction between water type and date being statistically significant (Table 2). The interaction between date and infection was also highly significant (Table 2).

This reflects the observation that, after the cystacanth stage being reached, the refuge score became weaker for infected individuals, while during the acanthella stage, the tendency was that infected individuals used more refuges than uninfected ones (Figure 2).

This observation is reinforced by the analysis of the effect sizes, but here differences appeared between scented and control water. Indeed, at acanthella stage, under non-scented water, the difference in refuge scores between infected and uninfected animals was always close to zero, indicating negligible effect of the infection (Figure 3). Under scented water, however, the values of δ were often between -0.11 and -0.4, i.e. a range where effect sizes are considered as small to medium (Vargha & Delaney, 2000; Kraemer & Kupfer, 2006). Nevertheless, only one value was significantly different from zero (95% C.I. did not overlap with zero), at day 55, i.e. at the late acanthella stage. After the cystacanth stage was reached, and here whatever the type of water, the effect size reversed, becoming positive, and can be considered as large, with values of δ exceeding 0.4 (Figure 3).

**Figure 3:**
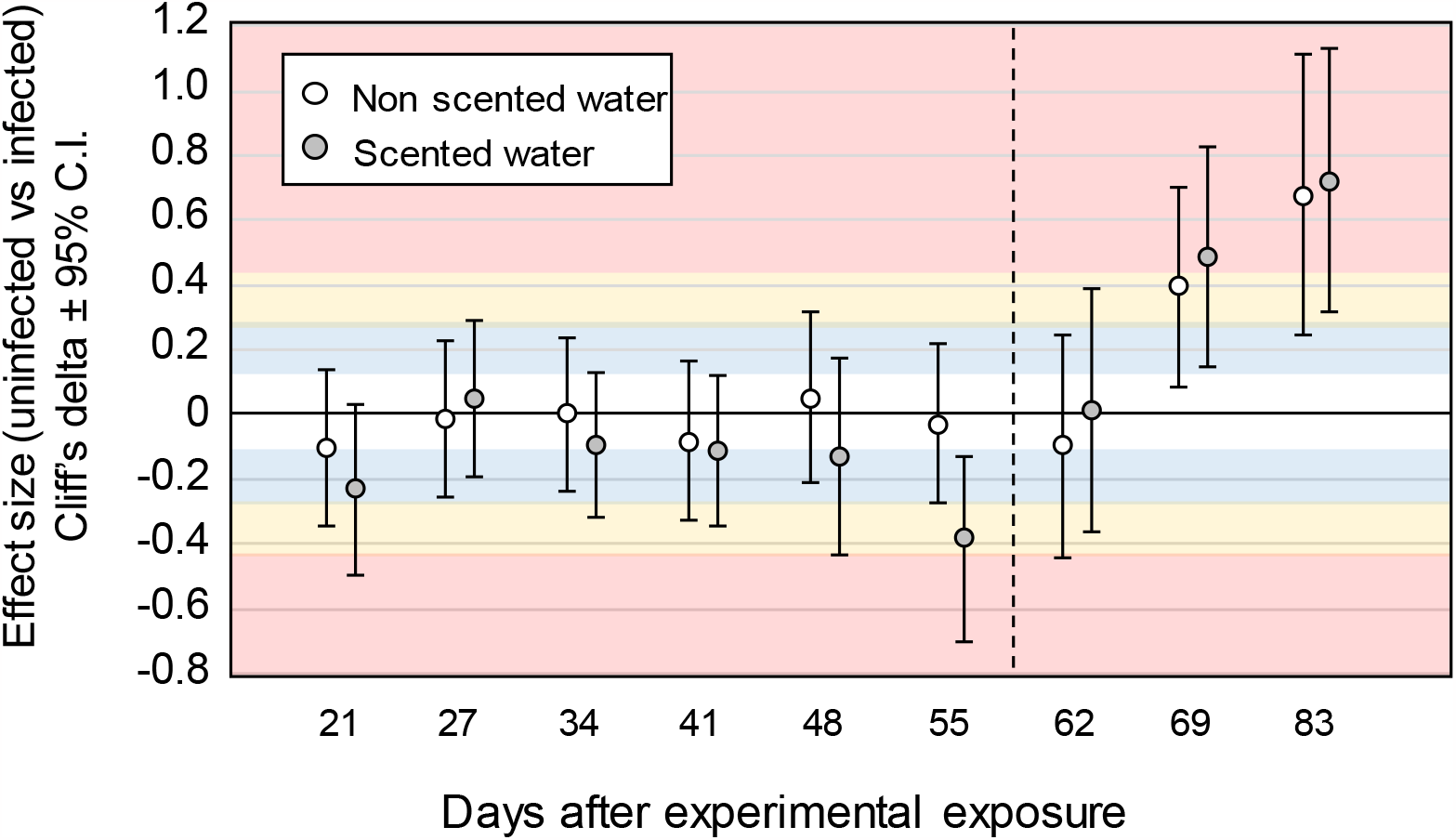
Effect sizes of refuge use of *Gammarus pulex* infected with *Pomphorhynchus laevis* compared to uninfected individuals, according to the date when the tests were made, separated by the water type in which they were tested. Points represent Cliff’s δ and error bars represent 95% confidence intervals. Positive values mean that the uninfected individuals used the refuge more than infected individuals, and negative values mean a lower use. The white, blue, yellow and red colors corresponded to areas where the effect size are considered as trivial, small, medium and large, respectively, using the comparisons between Cliff’s δ and Cohen’s d effect size made by (Kraemer & Kupfer, 2006). These areas must be seen as indicative. The dotted line denotes the transition between acanthella and cystacanth stages.

The effect of infection intensity was explored in infected animals only for the two dates where the analysis above suggested reversed effects of the infection, and where sample size was high enough to guarantee reasonable power, i.e. at days 55 and 69 post-exposure. At day 55, no effect of parasite number was seen on refuge use, nor on day 69. Indeed, after categorization as “1 parasite” and “more than 1 parasite”, Wilcoxon tests were not significant (z = -0.8144, p = 0.4097 and z = -1.3688, p = 0.1675, respectively). The same result was found if parasite number was treated as a discrete numerical variable since the correlations between parasite number and refuge use scores were non-significant for the two dates (Spearman’s ρ = 0.0595, p = 0.6379, n = 65 and Spearman’s ρ = 0.2333, p = 0.1229, n = 45, respectively).

The analysis of levels of activity revealed an effect of water type close to statistical significance, with significantly more time spent inactive in infected gammarids compared to uninfected ones, but also a highly significant increased inactivity with time (Figure 4, Table 3). None of the interactions were significant (Table 3).

**Figure 4:**
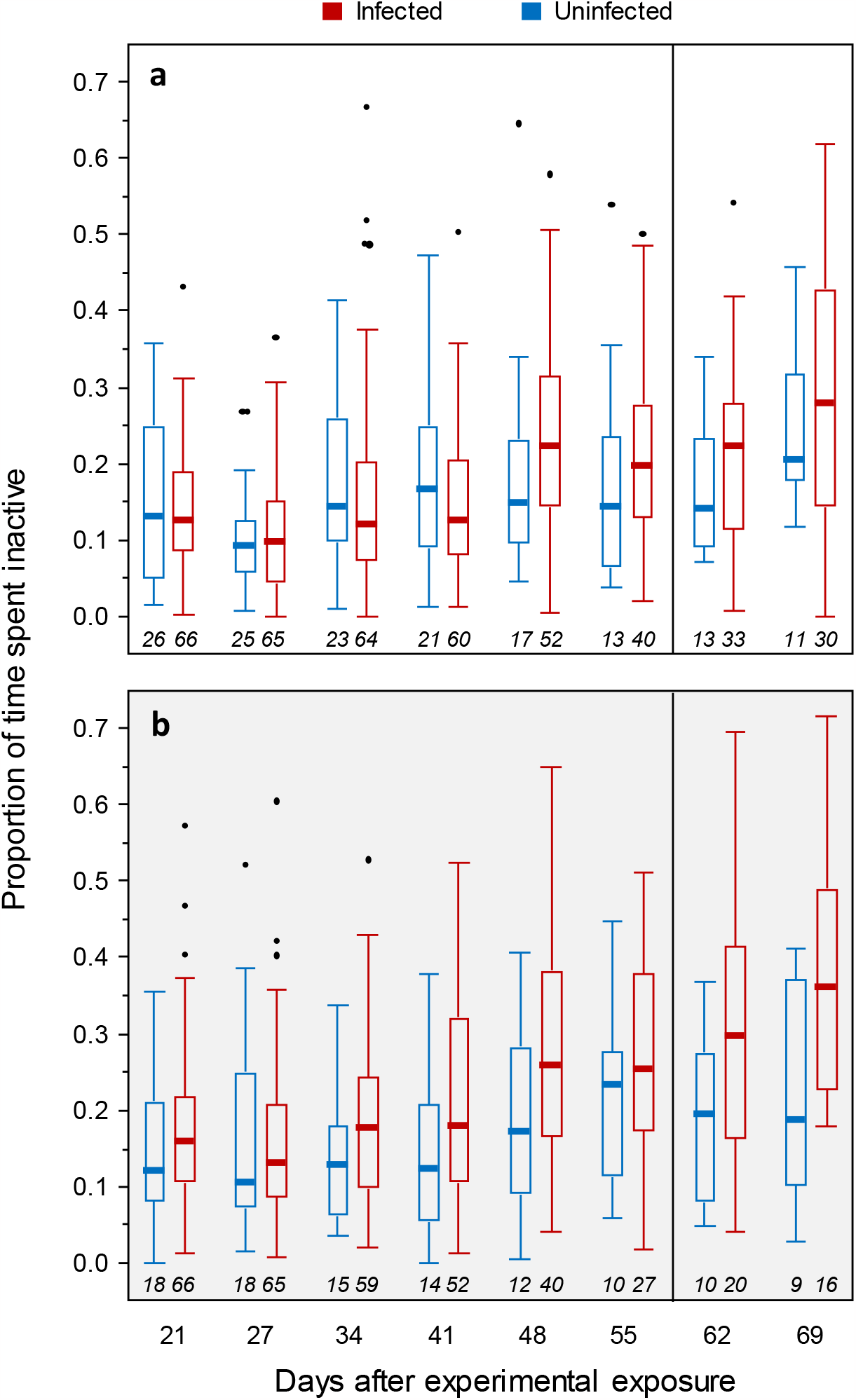
Proportion of time spent inactive in gammarids, according to their infection status by *Pomphorhynchus laevis*, between day 21 and day 69 post experimental exposure to parasite eggs. Gammarids were tested for 5 minutes in control water (a), or water signed with fish scent (b). Thick lines are the medians, boxes are the upper and lower quartiles, whiskers are the upper and lower 1.5*interquartile range, and dots represent outliers. The vertical black line denotes the transition between acanthella and cystacanth stages. Sample sizes are given in italics below each plot. Infection increased the gammarid’s inactivity whatever the water type and the developmental stage.

**Table 3:** Results of the model from the nparLD analysis, testing for the effects of water type (scented vs. control), infection status (infected with *P. laevis* vs. uninfected) and date of measurement on the activity rate of *G. pulex* individuals.

|  | AT statistics | df | P-value |
| --- | --- | --- | --- |
| Water type | 3.3850 | 1 | 0.0658 |
| **Infection** | **6.1807** | **1** | **0.0129** |
| **date** | **17.5550** | **4.8886** | **< 0.00001** |
| Water type * Infection | 2.5109 | 1 | 0.1131 |
| Water type * date | 1.1737 | 4.8886 | 0.3193 |
| Infection * date | 1.1683 | 4.8886 | 0.3221 |
| Water type * Infection * date | 1.1111 | 4.8886 | 0.3517 |

Analysis of effect sizes (Figure 5) confirmed but visually refined the analysis, with infected individuals being significantly more inactive than uninfected animals in the scented water.

**Figure 5:**
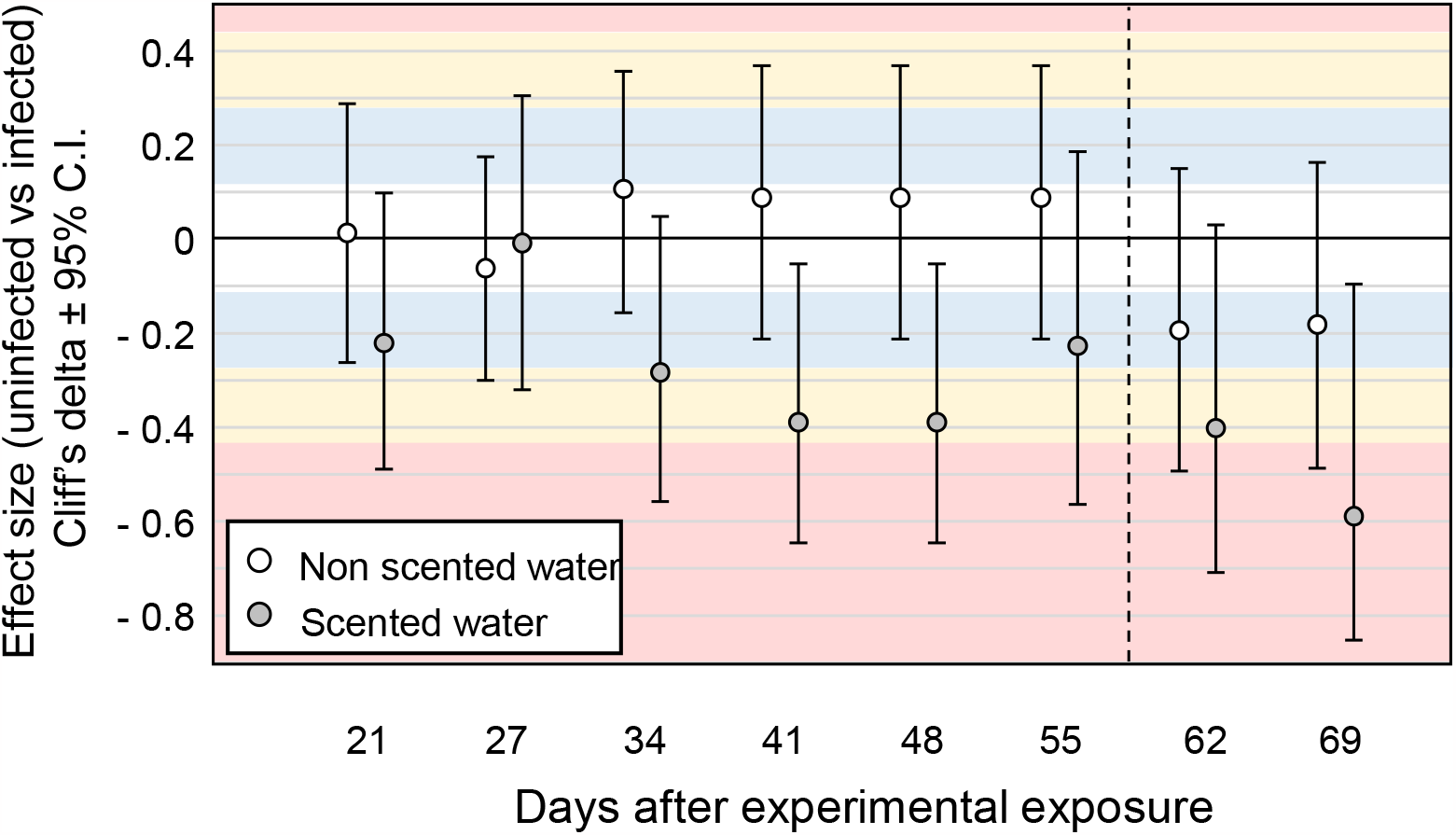
Effect sizes of time spent inactive in *Gammarus pulex* infected with *Pomphorhynchus laevis* compared to uninfected individuals, according to the dates when the tests were made and the water type. Points are Cliff’s δ and error bars represent 95% confidence intervals. Negative values mean that the infected individuals are more inactive than uninfected animals. The white, blue, yellow and red colors corresponded to areas where the effect size are considered as trivial, small, medium and large, respectively (see Figure 3). The dotted line denotes the transition between acanthella and cystacanth stages.

## Discussion

Our longitudinal survey of antipredatory behaviors in *Gammarus pulex* infected by *Pomphorhynchus laevis*, made in repeated measures, first confirmed previous observations that a shift occurred in refuge use between acanthella and cystacanth stages, i.e. between parasite stages non-infective and infective for the next host in the cycle (Dianne et al., 2011, 2014). This shift was nevertheless not observed for the activity rate, another behavior involved in predation avoidance (Andersson et al., 1986; Bollache et al., 2006). In addition, our study brings new information on this anti-predatory behavior.

First, confirming observations of Dianne et al. (2011), we found a difference in the ‘protective’ behavioral patterns of refuge use and activity rate according to the presence or absence of predatory cues. Indeed, while almost no difference between infected and uninfected animals was found when tests were made in control water, *G. pulex* infected by acanthellae use more the refuges at the end of their parasite development and are more inactive than uninfected ones in fish-scented water. Fish scent is definitely a source of stress for gammarids: it generally increases anti-predatory behavior when gammarids perceive predatory cues (e.g. they are repulsed by the chemical cues originating from fish (Perrot-Minnot et al., 2007) and reduce their activity (Andersson et al., 1986; Bollache et al., 2006). The infection by acanthellae would therefore strengthen this behavior. In another acanthocephalan parasite, *Polymorphus minutus*, the protective behavior has been observed in absence of predatory cues (Bailly et al., 2018). This parasite nevertheless shows a different nature of behavioral changes, probably in link with the difference in life cycle (*P. minutus* is using birds as definitive hosts), with probable different mechanisms involved (Tain et al., 2006; Perrot-Minnot et al., 2016).

The second information is that protective behaviors associated with the infection were significant – in scented water – at variable times during parasite ontogeny, depending on the behavior considered. This means that reinforcing the host’s antipredatory behavior outweighs the potential costs (Parker et al., 2009) differently depending on the behavior.

The reduced activity rate in infected animals, on the one hand, appeared quite early during acanthella growth (see a longer discussion on this trait in the next paragraph). The increase of refuge use, on the other hand, was found when acanthellae are growing old and was only significant at 55 days. Costs imposed by this behavioral change, as the one identified by Dianne et al. (2014), if imposed during a long time, may impact the survival of the host and may therefore limit in time the expression of protective manipulation during parasite ontogeny. In addition, for long-developing parasites, energy spent during growth may not be available for parasite manipulation (Poulin, 1994). It results that the moment when the expression of protection is the most optimal is when growth is complete, because energy then becomes available again. Therefore, the strategy of investing this energy in anti-predator behavior to safeguard past investments in growth can be selected. Interestingly, here, we observed that the time at which the protective component of manipulation is significant in *P. laevis* is when acanthella growth is complete, an observation compatible with such a strategy. Mechanistically, such a time course may be explained by chemicals secreted by parasites, that have the potential to be involved in host behavioral change (Berger et al., 2021, and references therein). However, we observed some variation (in scented water) on the differences between infected and uninfected gammarids: the effect size fluctuates between negligible values to medium values. This variation may be due to the fact that even uninfected gammarids did not use refuges constantly through time, a phenomenon that may interfere with the parasite’s consequences on host behavior. Indeed, from day 21 to day 55, there was a general trend of *G. pulex* using refuge more intensively as time is passing, and this independently from their infection status. After four behavioral measurements, all gammarids spent most of their time under the refuge, and this phenomenon is more intense in scented water. Gammarids, like other crustaceans are sensitive to stress, and one response to stress is to increase the intensity of anti-predatory behaviors (Fossat et al., 2014; Perrot-Minnot et al., 2017). Regular handling of shrimps increase their stress and modify the animal’s behavior (Takahashi, 2022). Therefore, we may infer that our experimental procedure was a source of stress for gammarids. Indeed, individuals were isolated for several weeks (gammarids are gregarious animals and isolation have consequences on their behavior, e.g. (Labaude et al., 2017) and were regularly handled to take them out of their isolation box and transfer them to a new environment (the aquarium or petri dish for behavioral tests). The observation that this phenomenon is more rapid under predatory cues is not surprising since predators are a major source of stress.

Whatever its mechanistic basis, this general increase in refuge use interferes with our ability to observe behavioral differences between gammarids infected or not by acanthellae, because it becomes less easy to find a difference in refuge use when this usage is already intense. After the parasite reached the cystacanth stage, we observed the shift in refuge use, and infected *G. pulex* are using the refuge less intensively than uninfected animals, as repeatedly observed in cases of infection with *P. laevis* (Franceschi, Bollache, et al., 2010; Dianne et al., 2011; Bauer & Rigaud, 2015; Fayard et al., 2020). The effect size of this exposure manipulation appeared higher than the one of protective manipulation in the present experiment, which was not the case in Dianne et al. (2011). However, in this later study, only one time point at acanthella and cystacanth stages each were measured, and the acanthella time-point was the one closest to cystacanth stage. If we compare only the absolute values of effect size at 55 and 69 days in our study, in the scented water, we will find no clear difference. However, by considering the whole ontogeny, it seems that the effect size of manipulation is greater at cystacanth stage than acanthella stage. Poulin (2010) noted that the relationship between the intensity of behavioral manipulation and the increase of predation rate (supposed to be due to manipulation) was not always straightforward, so we cannot really conclude about the efficiency of protection and exposure in our parasite-host system. However, since increase in mortality was mainly observed after the cystacanth stage being reached (as also observed by Labaude et al., 2020), it would make sense that the exposure manipulation is stronger than the protective manipulation because the selective pressure of reaching the definitive host is strong at that stage.

The third information of our study is that, as predicted, a second antipredatory behavior, namely the decreased activity rate, was also strengthened by infection during acanthella stage. Here again the differences in activity rate between infected and uninfected hosts were found mostly under predatory threat. This behavioural change appeared earlier in ontogeny than refuge use, suggesting that, if this change in behavior is adaptive for parasite, the costs of reducing activity are lower than increasing refuge use (Parker et al., 2009).

Indeed, in gammarids, a reduced activity is associated with limited metabolic expenditures rather than with costs (e.g. Normant et al., 2007), while increased refuge use induce costs on food intake (Dianne et al. 2014). However, and contrary to predictions, no shift in behavioral modification was observed for this behavior between acanthella and cystacanth stages, as the one observed for refuge use, the infected gammarids even being less active at cystacanth stages than at acanthella stages. Two alternative and exclusive hypotheses may be proposed for this observation. The first one is that the decreased activity rate in infected animals is a non-adaptive pathogenic byproduct of the infection, as sometimes suggested for some behavioral changes induced by parasites (Levri, 1999; Cézilly et al., 2010). Under this hypothesis, the progressive decrease according to time could just be interpreted as the long time spent by parasite in the host, decreasing progressively its mobility capacity. This could even be considered as a host strategy since here the interests of the parasite and the host are aligned during acanthella stage (reduced activity reduce predation risk, which is advantageous for both the parasite and the host, and save energy, which is advantageous for the host). Such hypotheses nevertheless explain neither why such a byproduct of the infection is not observed during acanthella stage in control water, nor the fact that the host and parasite interests became unaligned after the cystacanth stage is reached. Indeed, here, the reduced risk of predation due to low activity is advantageous for the host, but not for the parasite. The second hypothesis would be to consider that decreasing activity is adaptive for the parasite at both stages of development. During acanthella stage, this low activity rate would make the gammarid less conspicuous towards predators, as classically interpreted (Williams & Moore, 1985; Bollache et al., 2006). Notably, high rate of activity allows gammarids to initiate swimming and consequently enter the river current (drift) where predators are present (Williams & Moore, 1985; Schaffer et al., 2013). During the cystacanth stage, the animal’s tendency to search light (Franceschi et al., 2008) and avoid refuges (this study, Dianne et al., 2011) would already bring them into the river drift (McCahon et al., 1991; Lagrue et al., 2007). A low level of activity would therefore accentuate the predation exposure by reducing the swimming capacity of gammarids, which could be easily transported passively by the current of the river. Indeed, Lagrue et al. (2007) observed that the presence of *G. pulex* in the river drift is increased by 25-30 times when infected by *P. laevis*, the very same proportion of gammarids being found in the stomach of Bullhead predators. We could have here an example of parasite’s manipulation of a single host behavior that could be beneficial at all parasite stages, a characteristic that could have facilitated its selection.

To conclude, the first component of the ‘protective’ behavioral manipulation by the acanthocephalan *Pomphorhynchus laevis* in its host *Gammarus pulex*, namely the increased refuge use, was significant in late acanthella stage, only when predator cues were present in the environment. This antipredator behavior enhancement was not significant early in the *P. laevis* ontogeny, perhaps because of costs due to the long development time of parasites.

The refuge use then became weaker when the cystacanth stage was reached, illustrating the ‘exposure’ component of this parasitic manipulation. Another anti-predator behavior, the decrease in activity rate, was also expressed more strongly after the parasite infection, but showed no reversal during parasite ontogeny as observed for use of refuges. Differences in the intensity and direction of these two behavioral changes may be due to differences in their benefits relative to the developmental stage of the parasite: reduced host activity may be beneficial for transmission of the parasite to all parasite stages, while high and low refuge use are beneficial to the acanthella and cystacanth stages, respectively.

## Supporting information

supplementary files for Rigaud et al

## Funding

This study was funded by the French Agence Nationale de la Recherche (ANR), ANR-13-BSV7-0004-01

## Conflict of interest disclosure

The authors declare they have no financial conflicts of interest in relation to the content of the article.

## Author contributions

TR: Funding acquisition, Conceptualization, Experiments, Data curation, Statistical analysis, Writing – original draft, Writing – review and editing; ABal: Experiments, Data curation, Writing – review and editing; ABau: Statistical analysis, Writing – review and editing.

## Data access

https://doi.org/10.57745/AKA33Z

## Supplementary information

found at https://doi.org/10.1101/2023.04.25.538244. Includes comparisons between unexposed and uninfected-exposed animals and R codes used for analyses.

## Notes

### Competing Interest Statement

The authors have declared no competing interest.

### Summary of Updates

The revised version include the statement of a recommandation of this article by PCI Zoology on the tittle page.

## References

Abjornsson K, Hansson L-A, Brönmark C (2004) Responses of prey from habitats with different predator regimes: local adaptation and heritability. Ecology, 85, 1859–1866. 10.1890/03-0074

Andersson K, Brönmark C, Herrmann J, Malmqvist B, Otto C, Sjörström P (1986) Presence of sculpins (Cottus gobio) reduces drift activity of Gammarus pulex (Amphipoda). Hydrobiologia, 133, 209–215. 10.1007/BF00005592

Bailly Y, Cézilly F, Rigaud T (2018) Stage-dependent behavioural changes but early castration induced by the acanthocephalan parasite Polymorphus minutus in its Gammarus pulex intermediate host. Parasitology, 145, 260–268. 10.1017/S0031182017001457

Bauer A, Rigaud T (2015) Identifying a key host in an acanthocephalan-amphipod system. Parasitology, 142, 1588–1594. 10.1017/S0031182015001067

Benesh DP (2019) Tapeworm manipulation of copepod behaviour: parasite genotype has a larger effect than host genotype. Biology Letters, 15, 20190495. 10.1098/rsbl.2019.0495

Berdoy M, Webster JP, Macdonald DW (2000) Fatal attraction in rats infected with Toxoplasma gondii. Proceedings of the Royal Society B: Biological Sciences, 267, 1591–1594.

Berger CS, Laroche J, Maaroufi H, Martin H, Moon K-M, Landry CR, Foster LJ, Aubin-Horth N (2021) The parasite Schistocephalus solidus secretes proteins with putative host manipulation functions. Parasites & Vectors, 14, 436. 10.1186/s13071-021-04933-w

Bethel WM, Holmes JC (1977) Increased vulnerability of amphipods to predation owing to altered behavior induced by larval acanthocephalans. Canadian Journal of Zoology, 55, 110–115. 10.1139/z77-013

Bollache L, Kaldonski N, Troussard J-P, Lagrue C, Rigaud T (2006) Spines and behaviour as defences against fish predators in an invasive freshwater amphipod. Animal Behaviour, 72, 627–633. 10.1016/j.anbehav.2005.11.020

Cézilly F, Thomas F, Médoc V, Perrot-Minnot M-J (2010) Host-manipulation by parasites with complex life cycles: adaptive or not? Trends in Parasitology, 26, 311–317. 10.1016/j.pt.2010.03.009

Cliff N (1993) Dominance Statistics: Ordinal Analyses to Answer Ordinal Questions. Psychological Bulletin, 113, 494–509. 10.1037/0033-2909.114.3.494

Dianne L, Perrot-Minnot M-J, Bauer A, Gaillard M, Léger E, Rigaud T (2011) Protection first then facilitation: a manipulative parasite modulates the vulnerability to predation of its intermediate host according to its own developmental stage. Evolution, 65, 2692–2698. 10.1111/j.1558-5646.2011.01330.x

Dianne L, Perrot-Minnot M-J, Bauer A, Guvenatam A, Rigaud T (2014) Parasite-induced alteration of plastic response to predation threat: increased refuge use but lower food intake in Gammarus pulex infected with the acanothocephalan Pomphorhynchus laevis. International Journal for Parasitology, 44, 211–216. 10.1016/j.ijpara.2013.11.001

Fayard M, Dechaume-Moncharmont F, Wattier R, Perrot-Minnot M (2020) Magnitude and direction of parasite-induced phenotypic alterations: a meta-analysis in acanthocephalans. Biological Reviews, brv.12606. 10.1111/brv.12606

Fossat P, Bacqué-Cazenave J, De Deurwaerdère P, Delbecque J-P, Cattaert D (2014) Anxiety-like behavior in crayfish is controlled by serotonin. Science, 344, 1293–1297. 10.1126/science.1248811

Franceschi N, Bauer A, Bollache L, Rigaud T (2008) The effects of parasite age and intensity on variability in acanthocephalan-induced behavioural manipulation. International Journal for Parasitology, 38, 1161–1170. 10.1016/j.ijpara.2008.01.003

Franceschi N, Bollache L, Cornet S, Bauer A, Motreuil S, Rigaud T (2010) Co-variation between the intensity of behavioural manipulation and parasite development time in an acanthocephalan-amphipod system: Co-variation between the intensity of behavioural manipulation and parasite development. Journal of Evolutionary Biology, 23, 2143–2150. 10.1111/j.1420-9101.2010.02076.x

Franceschi N, Cornet S, Bollache L, Dechaume-Moncharmont F-X, Bauer A, Motreuil S, Rigaud T (2010) Variation between populations and local adaptation in acanthocephalan-induced parasite manipulation: variation in parasite-induced behavioral manipulation. Evolution, no-no. 10.1111/j.1558-5646.2010.01006.x

Hafer N (2018) Differences between populations in host manipulation by the tapeworm Schistocephalus solidus – is there local adaptation? Parasitology, 145, 762–769. 10.1017/S0031182017001792

Hafer-Hahmann N (2019) Experimental evolution of parasitic host manipulation. Proceedings of the Royal Society B: Biological Sciences, 286, 20182413. 10.1098/rspb.2018.2413

Hammerschmidt K, Koch K, Milinski M, Chubb JC, Parker GA (2009) When to go: optimization of host switching in parasites with complex life cycles. Evolution, 63, 1976–1986. 10.1111/j.1558-5646.2009.00687.x

Hughes DP, Brodeur J, Thomas F, UK P by RD University of Oxford (Eds.) (2012) Host Manipulation by Parasites. Oxford University Press, Oxford, New York.

Jensen, C. H., Weidner, J., Giske, J.,Jørgensen, C., Eliassen, S., & Mennerat, A. (2023). Adaptive host responses to infection can resemble parasitic manipulation. Ecology and Evolution, 13, e10318. 10.1002/ece3.10318

Kraemer HC, Kupfer DJ (2006) Size of Treatment Effects and Their Importance to Clinical Research and Practice. Biological Psychiatry, 59, 990–996. 10.1016/j.biopsych.2005.09.014

Labaude S, Cézilly F, De Marco L, Rigaud T (2020) Increased temperature has no consequence for behavioral manipulation despite effects on both partners in the interaction between a crustacean host and a manipulative parasite. Scientific Reports, 10, 11670. 10.1038/s41598-020-68577-z

Labaude S, Cézilly F, Tercier X, Rigaud T (2015) Influence of host nutritional condition on postinfection traits in the association between the manipulative acanthocephalan Pomphorhynchus laevis and the amphipod Gammarus pulex. Parasites & Vectors, 8, 403. 10.1186/s13071-015-1017-9

Labaude S, Rigaud T, Cézilly F (2017) Additive effects of temperature and infection with an acanthocephalan parasite on the shredding activity of Gammarus fossarum (Crustacea: Amphipoda): the importance of aggregative behavior. Global Change Biology, 23, 1415–1424. 10.1111/gcb.13490

Lagrue C, Kaldonski N, Perrot-Minnot MJ, Motreuil S, Bollache L (2007) Modification of hosts’ behavior by a parasite: field evidence for adaptive manipulation. Ecology, 88, 2839–2847. 10.1890/06-2105.1

Levri EP (1999) Parasite-induced change in host behavior of a freshwater snail: parasitic manipulation or byproduct of infection? Behavioral Ecology, 10, 234–241. 10.1093/beheco/10.3.234

McCahon CP, Maund SJ, Poulton MJ (1991) The effect of the acanthocephalan parasite (Pomphorhynchus laevis) on the drift of its intermediate host (Gammarus pulex). Freshwater Biology, 25, 507–513. 10.1111/j.1365-2427.1991.tb01393.x

Médoc V, Beisel J-N (2011) When trophically-transmitted parasites combine predation enhancement with predation suppression to optimize their transmission. Oikos, 120, 1452–1458. 10.1111/j.1600-0706.2011.19585.x

Moore J (1983) Responses of an Avian Predator and Its Isopod Prey to an Acanthocephalan Parasite. Ecology, 64, 1000–1015. 10.2307/1937807

Moore J (2002) Parasites and the Behavior of Animals. Oxford University Press.

Noguchi K, Gel YR, Brunner E, Konietschke F (2012) nparLD: An R Software Package for the Nonparametric Analysis of Longitudinal Data in Factorial Experiments. Journal of Statistical Software, 50, 1–23. 10.18637/jss.v050.i12

Normant M, Dziekonski M, Drzazgowski J, Lamprecht I (2007) Metabolic investigations of aquatic organisms with a new twin heat conduction calorimeter. Thermochimica Acta, 458, 101–106. 10.1016/j.tca.2007.01.025

Parker GA, Ball MA, Chubb JC, Hammerschmidt K, Milinski M (2009) When should a trophically transmitted parasite manipulate its host? Evolution, 63, 448–458. 10.1111/j.1558-5646.2008.00565.x

Perrot-Minnot M-J, Banchetry L, Cézilly F (2017) Anxiety-like behaviour increases safety from fish predation in an amphipod crustacea. Royal Society Open Science, 4, 171558. 10.1098/rsos.171558

Perrot-Minnot M-J, Kaldonski N, Cézilly F (2007) Increased susceptibility to predation and altered anti-predator behaviour in an acanthocephalan-infected amphipod. International Journal for Parasitology, 37, 645–651. 10.1016/j.ijpara.2006.12.005

Perrot-Minnot M, Maddaleno M, Cézilly F (2016) Parasite-induced inversion of geotaxis in a freshwater amphipod: a role for anaerobic metabolism? Functional Ecology, 30, 780–788. 10.1111/1365-2435.12516

Perrot-Minnot M-J, Sanchez-Thirion K, Cézilly F (2014) Multidimensionality in host manipulation mimicked by serotonin injection. Proceedings of the Royal Society B: Biological Sciences, 281, 20141915. 10.1098/rspb.2014.1915

Poulin R (1994) The evolution of parasite manipulation of host behaviour: a theoretical analysis. Parasitology, 109, S109–S118. doi:10.1017/s0031182000085127

Poulin R (2007) Evolutionary Ecology of Parasites: (Second Edition). Princeton University Press. 10.2307/j.ctt7sn0x

Poulin R (2010) Parasite Manipulation of Host Behavior. In: Advances in the Study of Behavior, pp. 151–186. Elsevier. 10.1016/S0065-3454(10)41005-0

Poulin R, Maure F (2015) Host manipulation by parasites: A look back before moving forward. Trends in Parasitology, 31, 563–570. 10.1016/j.pt.2015.07.002

Schaffer M, Winkelmann C, Hellmann C, Benndorf J (2013) Reduced drift activity of two benthic invertebrate species is mediated by infochemicals of benthic fish. Aquatic Ecology, 47, 99–107. 10.1007/s10452-013-9428-1

Svensson PA, Eghbal R, Eriksson R, Nilsson E (2022) How cunning is the puppet-master? Cestode-infected fish appear generally fearless. Parasitology Research, 121, 1305–1315. 10.1007/s00436-022-07470-2

Tain L, Perrot-Minnot M-J, Cézilly F (2006) Altered host behaviour and brain serotonergic activity caused by acanthocephalans: evidence for specificity. Proceedings of the Royal Society B: Biological Sciences, 273, 3039–3045. 10.1098/rspb.2006.3618

Takahashi K (2022) Changes in the anxiety-like and fearful behavior of shrimp following daily threatening experiences. Animal Cognition, 25, 319–327. 10.1007/s10071-021-01555-8

Vargha A, Delaney HD (2000) A Critique and Improvement of the CL Common Language Effect Size Statistics of McGraw and Wong. Journal of Educational and Behavioral Statistics, 25, 101–132.

Weinreich F, Benesh DP, Milinski M (2013) Suppression of predation on the intermediate host by two trophically-transmitted parasites when uninfective. Parasitology, 140, 129–135. 10.1017/S0031182012001266

Williams DD, Moore KA (1985) The Role of Semiochemicals in Benthic Community Relationships of the Lotic Amphipod Gammarus pseudolimnaeus: A Laboratory Analysis. Oikos, 44, 280. 10.2307/3544701

Wudkevich K, Wisenden BD, Chivers DP, Smith RJF (1997) Reactions of Gammarus lacustris to Chemical Stimuli from Natural Predators and Injured Conspecifics. Journal of Chemical Ecology, 23, 1163–1173. 10.1023/B:JOEC.0000006393.92013.36

