## supplementary files for Rigaud et al for "Time-course of antipredator behavioral changes induced by the helminth *Pomphorhynchus laevis* in its intermediate host *Gammarus pulex*: the switch in manipulation according to parasite developmental stage differs between behaviors"

### Supplementary material

##### Comparison between Unexposed and Uninfected-exposed animals for their survival

Exposed-uninfected animals did not differ significantly from unexposed gammarids for their survival (Figure S1), whatever the water type in which they were maintained (Table S1).

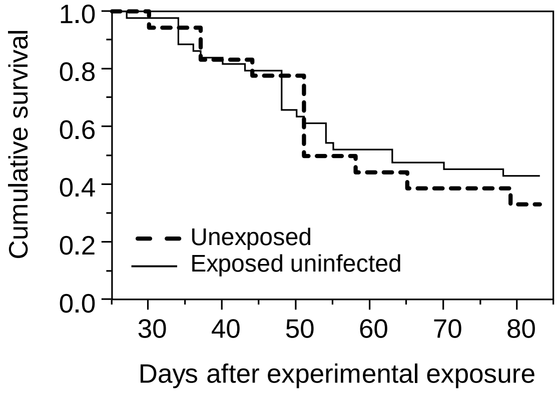


**Figure S1:** Comparison of cumulative survival between *Gammarus pulex* experimentally exposed to *Pomphorhynchus laevis*, but uninfected (n = 44), and control, unexposed, animals (n = 18).

**Table S1:** Comparison of survival between *Gammarus pulex* experimentally exposed to *Pomphorhynchus laevis*, but uninfected (n = 44), and control, unexposed, animals (n = 18), taking into account the water type in which they were maintained (fish-scented vs. control water).

| Source | d.f. | z | p |
|---|---|---|---|
| Water type | 1 | -0.516 | 0.606 |
| Exposure | 1 | -0.457 | 0.648 |
| Water*exposure | 1 | 0.070 | 0.944 |

Test of the proportional hazards assumption $\chi 2$ = 1.4192; 3 d.f.; p = 0.7

##### Refuge use among the three groups of *Gammarus pulex* : unexposed to *Pomphorynchus laevis*, exposed but not infected, exposed and infected

Refuge use according to these three groups are described on Figure S2 and analyzed in tables S2 to S4. Unexposed *G. pulex* were mistakenly not measured for their refuge use at day 83, (Figure S2). Therefore, the analyses taking into account unexposed animals were only run until day 69. However, the comparison between Exposed Infected and Exposed Uninfected animals was possible until day 83.

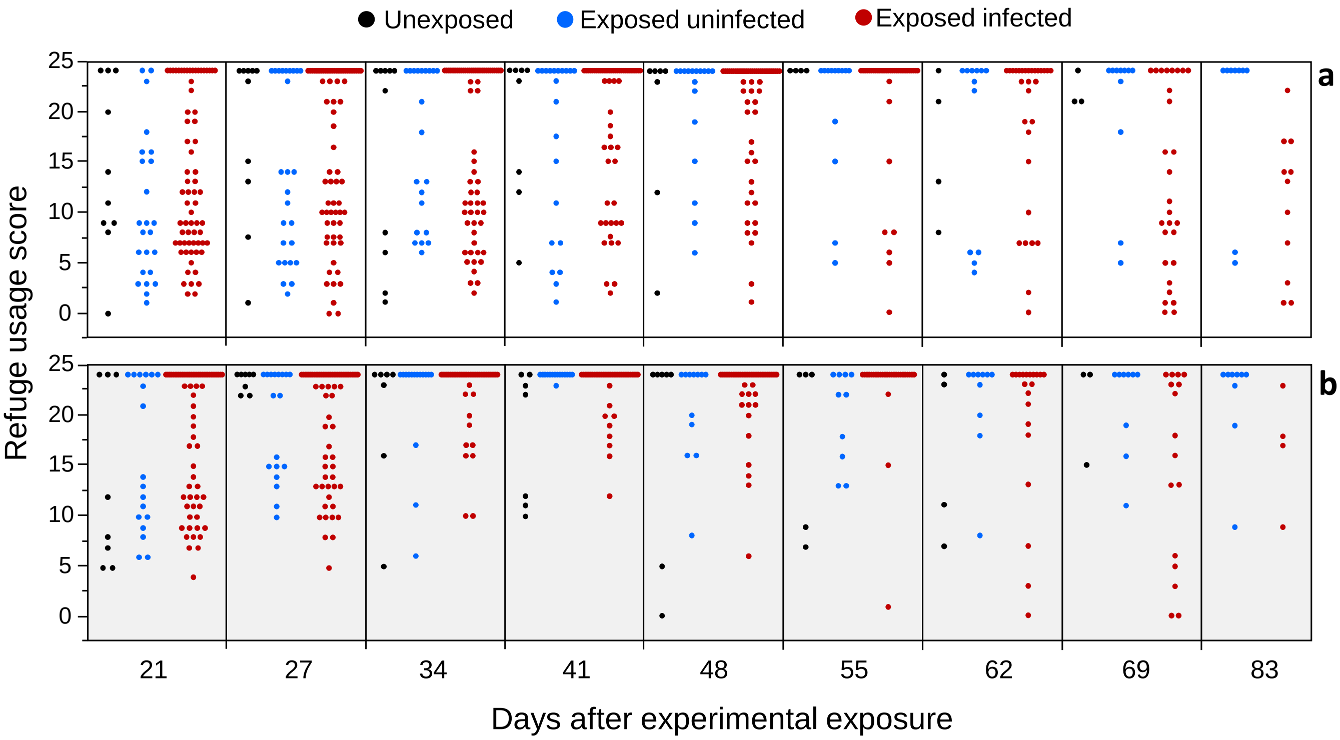


**Figure S2:** Scores of refuge use by gammarids, according to their exposure and infection status by *Pomphorhynchus laevis*, between day 21 and day 83 post-experimental exposure to parasite eggs. Gammarids were tested in control water (a), or water signed with fish scent (b). Scores range from 0 (individuals always outside the refuge) to 24 (individuals always inside the refuge). Each dot represent the score of a single individual.

The refuge use by gammarids varied considerably with time (Figure S2, Table S2), being most of the time moderate at the beginning of our survey, becoming progressively more intensive until reaching high values around 30 to 50 days. This phenomenon was found whatever the infection status and the water type, but was found to be more rapid under scented water than control water (Figure 2), the interaction between water type and date being statistically significant (Table S2). The interaction between date and infection was also highly significant (Table S2).

**Table S2:** Results of the model from the nparLD analysis, testing for the effects of water type (scented vs. control), infection status (Unexposed to *P. laevis*; exposed-uninfected; exposed-infected) and date of measurement on the scores of refuge use by *G. pulex* individuals. The analysis was only possible until day 69, because unexposed individuals were not available later.

| | AT statistics | df | P-value |
|---|---|---|---|
| **Water type** | **4.2856** | **1** | **0.0384** |
| Infection | 0.0265 | 1.6069 | 0.9518 |
| **date** | **9.4245** | **5.3362** | **< 0.00001** |
| Water type * Infection | 1.0180 | 1.6069 | 0.3475 |
| **Water type * date** | **2.4376** | **5.3362** | **0.0289** |

| | | | |
|---|---|---|---|
| **Infection * date** | **2.8393** | **7.8442** | **0.0040** |
| Water type * Infection * date | 0.9302 | 7.8442 | 0.4886 |

One week after the cystacanth stage being reached, the refuge score became weak for infected individuals, while during the acanthella stage, they used refuges at least as intensively as unexposed or exposed-uninfected ones (Figure S2).

If removing unexposed individuals from the analysis, because of their small sample size and their unavailability at day 83, the analysis remain essentially the same (Table S3).

**Table S3:** Results of the model from the nparLD analysis, testing for the effects of water type (scented vs. control), infection status (Exposed-uninfected; exposed-infected by *P. laevis*) and date of measurement on the scores of refuge use by *G. pulex* individuals. Here, the analysis was possible until day 83.

| | AT statistics | df | P-value |
|---|---|---|---|
| **Water type** | **10.1615** | **1** | **0.0014** |
| Infection | 0.5262 | 1 | 0.4682 |
| **date** | **13.8168** | **5.3410** | **< 0.00001** |
| Water type * Infection | 0.0468 | 1 | 0.8287 |
| **Water type * date** | **3.2153** | **5.3410** | **0.0054** |
| **Infection * date** | **10.4477** | **5.3410** | **< 0.00001** |
| Water type * Infection * date | 0.7463 | 5.3410 | 0.5972 |

Finally, the comparison on Unexposed individuals and Exposed-infected animals showed non-significant differences, except the date that still influence the animal's behaviour, but in the same way according to exposure status (Table S4).

**Table S4:** Results of the model from the nparLD analysis, testing for the effects of water type (scented vs. control), infection status (Unexposed to *P. laevis* vs. Exposed-uninfected) and date of measurement on the scores of refuge use by *G. pulex* individuals. The analysis was only possible until day 69, because unexposed individuals were not available later.

| | AT statistics | df | P-value |
|---|---|---|---|
| Water type | 1.5220 | 1 | 0.2173 |
| Infection | 1.7806 | 1 | 0.9966 |
| **date** | **5.5356** | **5.2070** | **<0.0001** |
| Water type * Infection | 1.3851 | 1 | 0.2392 |
| Water type * date | 1.6531 | 5.2070 | 0.3915 |
| Infection * date | 1.6090 | 5.2070 | 0.1508 |

| | | | |
|---|---|---|---|
| Water type * Infection * date | 0.8824 | 5.2070 | 0.4950 |

These results confirmed previous ones showing that hosts exposed but non-infected by *P. laevis* survive, behave and have a physiology similar to unexposed ones (Franceschi et al. 2008, Sanchez-Thirion et al. 2019, Cornet et al. 2009, respectively, and Bauer Unpublished results). Therefore, to increase sample size, notably in the three last dates, we cumulated Unexposed and Exposed-uninfected animals for comparing their refuge usage with that of infected animals. The results are similar to those exposed in table S2 (see main text).

**Scripts R for analyses**

```
######################################################################
### anova non paramétrique de mesures répétées avec données manquantes
######################################################################



### ATTENTION !
### Si le modèle donne cette erreur, c'est qu'un des facteurs n'est pas
mesuré jusqu'à la fin de la série temporelle :
### Erreur dans qr.default(V) : NA/NaN/Inf dans un appel à une fonction
externe (argument 1)



##########################################################
###  packages nécessaires
##########################################################

# install.packages("nparLD")
library(nparLD)
# help(nparLD)

# package pour récup le .xls
# install.packages("xlsx")
library(xlsx)

# pour calculer le d de cohen
# install.packages("effsize")
library(effsize)

# install.packages("tidyr")
library(tidyr)
# pour les fonctions replace_na ou drop_na




##########################################################
###  import du jeu de données et nettoyage
##########################################################

### lecture du fichier excel
brut <- read.xlsx2(file.choose(), sheetIndex=1, header=T,
colClasses="character")
```

```
donnee <- brut


### transformer les scores en variable numérique
donnee$score_refuge <- as.numeric(donnee$score_refuge)

summary(donnee)

table(donnee$type_eau, donnee$infection)


### enlever la première date
donnee <- donnee[donnee$jours != 0, ]
table(donnee$jours)





##########################################################
###  construction du modèle global
##########################################################


travail <- donnee

### enlever dernière date pour pouvoir comparer les témoins
travail <- travail[travail$jours != "83", ]
table(travail$jours)

table(travail$type_eau, travail$infection)

### assignation des variables principales
var <- travail[ ,"score_refuge"]
time <- travail[ ,"jours"]
subject <- travail[ ,"individu"]

### 2 variables explicatives => plan F2-LD-F1
group1 <- travail[ ,"type_eau"]
group2 <- travail[ ,"infection"]

### stat
modele <- f2.ld.f1(var, time, group1, group2, subject,
        time.name="date", group1.name="type eau", group2.name="statut
infectieux",
        description=FALSE)
#  Warning(s): The covariance matrix is singular.


modele$Wald.test

modele$ANOVA.test



### rep graphique

non_signee <- travail[travail$type_eau == "NS", ]
signee <- travail[travail$type_eau == "S", ]
```

```
boxplot(
        score_refuge ~ infection * jours, data = travail,
        axes = FALSE,
        ylim = c(0,40),
        xlab = "jours",
        ylab = "score 'photophilie'",
        col = c(1, 2, 3), lwd = 2
        )

box ()

axis(
        1,
        c(2, 5, 8, 11, 14, 17,20, 23), tick = FALSE,
        labels = c(21, 27, 34, 41, 48, 55, 62, 69),
        line=-1, cex.axis=0.8
        )

axis(side = 2, at = seq(from=0, to=1, by= 0.2), tck = -0.03, las = 1)




##########################################################
###  Comparaison sni et controles
##########################################################


### comparison controles et soumis non infectés

travail <- donnee

### enlever dernière date pour pouvoir comparer les témoins
travail <- travail[travail$jours != "83", ]
table(travail$jours)


### on enlève les parasités

travail <- travail[travail$infection != "p", ]
table(travail$infection)
table(travail$type_eau, travail$infection)


### assignation des variables principales
var <- travail[ ,"score_refuge"]
time <- travail[ ,"jours"]
subject <- travail[ ,"individu"]

### 2 variables explicatives => plan F2-LD-F1
group1 <- travail[ ,"type_eau"]
group2 <- travail[ ,"infection"]

### modèle
modele <- f2.ld.f1(var, time, group1, group2, subject,
        time.name="date", group1.name="type eau", group2.name="statut
infectieux", description=FALSE
        )
# Warning(s): The covariance matrix is singular.
```

```
modele$Wald.test

modele$ANOVA.test





##########################################################
###  Comparaison controles et para
##########################################################

travail <- donnee

### enlever dernière date pour pouvoir comparer les témoins
travail <- travail[travail$jours != "83", ]
table(travail$jours)

### on enlève les sni
travail <- travail[travail$infection != "eni", ]
table(travail$infection)
table(travail$type_eau, travail$infection)

### assignation des variables principales
var <- travail[ ,"score_refuge"]
time <- travail[ ,"jours"]
subject <- travail[ ,"individu"]

### 2 variables explicatives => plan F2-LD-F1
group1 <- travail[ ,"type_eau"]
group2 <- travail[ ,"infection"]

### modèle
modele <- f2.ld.f1(var, time, group1, group2, subject,
        time.name="date", group1.name="type eau", group2.name="statut
infectieux", description=FALSE
        )
# Warning(s): The covariance matrix is singular.


modele$Wald.test

modele$ANOVA.test





##########################################################
###  construction du modèle comparaison sni et para
##########################################################

travail <- donnee

### on enlève les controles
travail <- travail[travail$infection != "control", ]
table(travail$infection)
table(travail$type_eau, travail$infection)

### assignation des variables principales
```

```
var <- travail[ ,"score_refuge"]
time <- travail[ ,"jours"]
subject <- travail[ ,"individu"]

### 2 variables explicatives => plan F2-LD-F1
group1 <- travail[ ,"type_eau"]
group2 <- travail[ ,"infection"]

### modèle
modele <- f2.ld.f1(var, time, group1, group2, subject,
        time.name="date", group1.name="type eau", group2.name="statut
infectieux", description=FALSE
        )
# pas de warning


modele$Wald.test

modele$ANOVA.test


-----------------------------------------------------------------------------------------------------------------


##########################################################
# survie en fonction du type d'eau et du statut infectieux,
# comparaisons gammares témoins, soumis non infectés et infections labo
##########################################################





##########################################################
###  packages nécessaires
##########################################################

# package pour récup le .xls
# install.packages("xlsx")
library(xlsx)


# pour les régressions de cox et les graphes de survie
# install.packages('survival')
library(survival)


# pour la fonction replace_na si nécessaire
# install.packages("tidyr")
library(tidyr)





##########################################################
###  import data
##########################################################

### récup fichier xlsx
```

```
brut <- read.xlsx2(file.choose(), sheetIndex=1, header=T,
colClasses="character")




##########################################################
###  nettoyage
##########################################################

donnee <- brut
nrow(donnee)

str(donnee)

donnee$evenement <- as.numeric(donnee$evenement)
donnee$duree <- as.numeric(donnee$duree)




##########################################################
###  Survie gammares non infectés :
###  sni / témoins
##########################################################

### ---------------------------------------------------
###  selection données, effectifs
### ---------------------------------------------------

travail <- donnee

### recodage statut infectieux
travail <- travail[travail$infection != "p", ]
travail$lot_def[travail$infection == "n"] <- "exposed uninfected"
travail$lot_def[travail$infection == "T"] <- "control"

table(travail$type_eau, travail$lot_def)



### ---------------------------------------------------
###  régressions
### ---------------------------------------------------

reg <- coxph(Surv(duree, evenement) ~ type_eau + lot_def +
type_eau*lot_def, data = travail)
summary(reg)

### test proportionnalité des risques
cox.zph(reg)
plot(cox.zph(reg))



### ---------------------------------------------------
###  courbes de survie
### ---------------------------------------------------
```

```
# paramètres graphiques
#dev.new(width = 6, height = 3)
#par(mfrow = c(1,2),
#       mar = c(4,4,1,1),
#       cex=1
#       )


### infos pour graphique
abscisse <- c(0, 90)
# traits = 1
couleur <- c(4, 2)
legende <- c("control water", "scented water", "control", "exposed not
infected")
couleur_legende <- c(1, 1, 4, 2)

###
eau_signee <- travail[travail$type_eau == "S", ]
eau_non_signee <- travail[travail$type_eau == "NS", ]

###
plot(
               survfit(Surv(duree, evenement) ~ lot_def, data =
eau_non_signee),
               lwd = 2,
               lty = 1,
               col = couleur,
               cex.axis = 1,
               las = 1,
               tck = -0.03,
               xlim = abscisse
       )

par(new=TRUE)

plot(
               survfit(Surv(duree, evenement) ~ lot_def, data =
eau_signee),
               lwd = 2,
               lty = 2,
               col = couleur,
               cex.axis = 1,
               las = 1,
               tck = -0.03,
               xlim = abscisse
       )

legend(
               "bottomleft",
               legend = legende,
               col = couleur_legende,
               #lty = traits, ne pas préciser pour l'export ppt
               bty = "n",
               cex = 0.8,
               lty = c(1,2,1,1),
               lwd = 2,
               text.col = couleur_legende,
               horiz = FALSE,
               inset = c(0.01, 0.01)
       )
```

```
#########################################################
###  Survie en fonction du statut infectieux :
###  parasité / non parasité (témoins + sni)
#########################################################

### ----------------------------------------------------
###  selection données, effectifs
### ----------------------------------------------------

travail <- donnee

### recodage statut infectieux
travail$lot_def[travail$infection == "p"] <- "infected"
travail$lot_def[travail$infection == "n"] <- "uninfected"
travail$lot_def[travail$infection == "T"] <- "uninfected"

table(travail$type_eau, travail$lot_def)



### ----------------------------------------------------
###  régressions
### ----------------------------------------------------

reg <- coxph(Surv(duree, evenement) ~ type_eau + lot_def +
type_eau*lot_def, data = travail)
summary(reg)
cox.zph(reg)
plot(cox.zph(reg))


reg <- coxph(Surv(duree, evenement) ~ type_eau , data = travail)
summary(reg)
cox.zph(reg)
plot(cox.zph(reg))



### sans le facteur 'eau'
reg <- coxph(Surv(duree, evenement) ~ lot_def , data = travail)



### ----------------------------------------------------
###  courbes de survie
### ----------------------------------------------------

# infos pour graphique
abscisse <- c(0, 90)
# traits = 1
couleur <- c(2, 4)
legende <- c("control water", "scented water", "infected", "not infected")
couleur_legende <- c(1, 1, 2, 4)

###
eau_signee <- travail[travail$type_eau == "S", ]
eau_non_signee <- travail[travail$type_eau == "NS", ]
```

```
###
plot(
		survfit(Surv(duree, evenement) ~ lot_def, data =
eau_non_signee),
		lwd=2,
		lty = 1,
		col = couleur,
		cex.axis=1,
		las=1,
		tck =-0.03,
		xlim = c(0,90)
	)

par(new=TRUE)

plot(
		survfit(Surv(duree, evenement) ~ lot_def, data =
eau_signee),
		lwd=2,
		lty = 2,
		col = couleur,
		cex.axis=1,
		las=1,
		tck =-0.03,
		xlim = c(0,90)
	)

legend(
		"bottomleft",
		legend = legende,
		col = couleur_legende,
		#lty = traits, ne pas préciser pour l'export ppt
		bty = "n",
		cex = 0.8,
		lty = c(1,2,1,1),
		lwd = 2,
		text.col = couleur_legende,
		horiz = FALSE,
		inset = c(0.01, 0.01)
	)




##########################################################
###  Survie période acanthelles
###  entre 0 et 59 jours
##########################################################



### ---------------------------------------------------
###  selection données, effectifs
### ---------------------------------------------------

travail <- donnee

### censurer à 55 jours
for (i in 1: nrow(travail)) {
```

```
        if(travail$duree[i] > 59) {
                travail$evenement[i] <- 0
                }
        }
travail$duree[travail$duree > 59] <- 59
table(travail$duree)

### recodage statut infectieux
travail$lot_def[travail$infection == "n"] <- "uninfected"
travail$lot_def[travail$infection == "T"] <- "uninfected"
travail$lot_def[travail$infection == "p"] <- "infected"

table(travail$type_eau, travail$lot_def)



### --------------------------------------------------
###  régressions
### --------------------------------------------------

reg <- coxph(Surv(duree, evenement) ~ type_eau + lot_def +
type_eau*lot_def, data = travail)

summary(reg)
# vraiment pas d'effet du statut infectieux ni de l'interaction ; type_eau
limite

cox.zph(reg)
plot(cox.zph(reg))

### on teste juste le type d'eau
reg <- coxph(Surv(duree, evenement) ~ type_eau, data = travail)
# NS



### --------------------------------------------------
###  courbes de survie
### --------------------------------------------------

### infos pour graphique
abscisse <- c(20, 60)
# traits = 1
couleur <- c(2, 4)
legende <- c("control water", "scented water", "infected", "not infected")
couleur_legende <- c(1, 1, 2, 4)

### séparer par type d'eau
eau_signee <- travail[travail$type_eau == "S", ]
eau_non_signee <- travail[travail$type_eau == "NS", ]

### graphique
plot(
                survfit(Surv(duree, evenement) ~ lot_def, data =
eau_non_signee),
                lwd = 2,
                lty = 1,
                col = couleur,
                cex.axis = 1,
                las = 1,
                tck = -0.03,
```

```
                xlim = abscisse
        )

par(new=TRUE)

plot(
                survfit(Surv(duree, evenement) ~ lot_def, data =
eau_signee),
                lwd = 2,
                lty = 1,                      # lty=2 ; mettre à 1 pour
export graphique vecto
                col = couleur,
                cex.axis = 1,
                las = 1,
                tck = -0.03,
                xlim = abscisse
        )

legend(
                "bottomleft",
                legend = legende,
                col = couleur_legende,
                #lty = traits, ne pas préciser pour l'export ppt
                bty = "n",
                cex = 0.8,
                lty = c(1,2,1,1),
                lwd = 2,
                text.col = couleur_legende,
                horiz = FALSE,
                inset = c(0.01, 0.01)
        )




###########################################################
###  Survie période cystacanthes à partir de 56 jours :
###  début quand les parasites sont encore tous au stade acanthelle
###########################################################

### ---------------------------------------------------
###  selection données, effectifs
### ---------------------------------------------------

travail <- donnee

### enlever les gammares morts avant 56 jours
travail <- travail[travail$duree >= 56, ]
table(travail$duree)

### recoder statut infectieux
travail$lot_def[travail$infection == "n"] <- "uninfected"
travail$lot_def[travail$infection == "T"] <- "uninfected"
travail$lot_def[travail$infection == "p"] <- "infected"

table(travail$type_eau, travail$lot_def)



### ---------------------------------------------------
```

```
###  régressions
### ---------------------------------------------------


reg <- coxph(Surv(duree, evenement) ~ type_eau + lot_def +
type_eau*lot_def, data = travail)

summary(reg)
# seulement effet du statut infectieux

cox.zph(reg)
plot(cox.zph(reg))
# OK




### ---------------------------------------------------
###  courbes de survie
### ---------------------------------------------------

# infos pour graphique

abscisse <- c(55, 83)
# traits = 1
couleur <- c(2, 4)
legende <- c("control water", "scented water", "infected", "not infected")
couleur_legende <- c(1, 1, 2, 4)

eau_signee <- travail[travail$type_eau == "S", ]
eau_non_signee <- travail[travail$type_eau == "NS", ]

plot(
               survfit(Surv(duree, evenement) ~ lot_def, data =
eau_non_signee),
               lwd = 2,
               lty = 1,
               col = couleur,
               cex.axis = 1,
               las = 1,
               tck = -0.03,
               xlim = abscisse,
               ylim = c(0,1)
        )

par(new=TRUE)

plot(
               survfit(Surv(duree, evenement) ~ lot_def, data =
eau_signee),
               lwd = 2,
               lty = 2,
               col = couleur,
               cex.axis = 1,
               las = 1,
               tck = -0.03,
               xlim = abscisse,
               ylim = c(0,1)
        )

legend(
               "bottomleft",
```

```
               legend = legende,
               col = couleur_legende,
               #lty = traits, ne pas préciser pour l'export ppt
               bty = "n",
               cex = 0.8,
               lty = c(1,2,1,1),
               lwd = 2,
               text.col = couleur_legende,
               horiz = FALSE,
               inset = c(0.01, 0.01)
        )




##########################################################
###  Survie période cystacanthes, à partir de 60 jours :
###  une fois que tous les parasites sont des cystacanthes
##########################################################



### ---------------------------------------------------
###  selection données, effectifs
### ---------------------------------------------------

travail <- donnee

### enlever les gammares morts avant 60 jours
travail <- travail[travail$duree >= 60, ]
table(travail$duree)

### recoder statut infectieux
travail$lot_def[travail$infection == "n"] <- "uninfected"
travail$lot_def[travail$infection == "T"] <- "uninfected"
travail$lot_def[travail$infection == "p"] <- "infected"

table(travail$type_eau, travail$lot_def)



### ---------------------------------------------------
###  régressions
### ---------------------------------------------------

reg <- coxph(Surv(duree, evenement) ~ type_eau + lot_def +
type_eau*lot_def, data = travail)

summary(reg)
# seulement effet du statut infectieux

cox.zph(reg)
plot(cox.zph(reg))
# OK



### ---------------------------------------------------
###  courbes de survie
### ---------------------------------------------------
```

```
# infos pour graphique

abscisse <- c(60, 83)
# traits = 1
couleur <- c(2, 4)
legende <- c("control water", "scented water", "infected", "not infected")
couleur_legende <- c(1, 1, 2, 4)

eau_signee <- travail[travail$type_eau == "S", ]
eau_non_signee <- travail[travail$type_eau == "NS", ]

plot(
		survfit(Surv(duree, evenement) ~ lot_def, data =
eau_non_signee),
		lwd = 2,
		lty = 1,
		col = couleur,
		cex.axis = 1,
		las = 1,
		tck = -0.03,
		xlim = abscisse,
		ylim = c(0,1)
	)

par(new=TRUE)

plot(
		survfit(Surv(duree, evenement) ~ lot_def, data =
eau_signee),
		lwd = 2,
		lty = 2,
		col = couleur,
		cex.axis = 1,
		las = 1,
		tck = -0.03,
		xlim = abscisse,
		ylim = c(0,1)
	)

legend(
		"bottomleft",
		legend = legende,
		col = couleur_legende,
		#lty = traits, ne pas préciser pour l'export ppt
		bty = "n",
		cex = 0.8,
		lty = c(1,2,1,1),
		lwd = 2,
		text.col = couleur_legende,
		horiz = FALSE,
		inset = c(0.01, 0.01)
	)




##########################################################
###  effet de la charge parasitaire
##########################################################
```

```
travail <- donnee

travail$n.parasite <- as.numeric(travail$n.parasite)

infectes <- travail[travail$infection == "p", ]



### choisir le codage de l'intensité

table(infectes$n.parasite)
reg <- coxph(Surv(duree, evenement) ~ n.parasite, data = infectes) #
continu
summary(reg)

table(infectes$n.para.cat)
reg <- coxph(Surv(duree, evenement) ~ n.para.cat, data = infectes) # 1, 2,
>2
summary(reg)

table(infectes$n.para.cat2)
reg <- coxph(Surv(duree, evenement) ~ n.para.cat2, data = infectes) # 1, >1
summary(reg)



### avec le facteur 'type d'eau'

table(infectes$n.para.cat2, infectes$type_eau)
reg <- coxph(Surv(duree, evenement) ~ n.para.cat2 + type_eau +
n.para.cat2*type_eau, data = infectes) # 1, >1
summary(reg)

cox.zph(reg)




# fin propre ---------------------------------------------------
```
